# A continuum membrane model predicts curvature sensing by helix insertion

**DOI:** 10.1101/2021.04.22.440963

**Authors:** Yiben Fu, Wade F. Zeno, Jeanne C. Stachowiak, Margaret E. Johnson

## Abstract

Protein domains, such as ENTH (Epsin N-terminal homology) and BAR (bin/amphiphysin/rvs), contain amphipathic helices that drive preferential binding to curved membranes. However, predicting how the physical parameters of these domains control this ‘curvature sensing’ behavior is challenging due to the local membrane deformations generated by the nanoscopic helix on the surface of a large sphere. To overcome this challenge, we here use a deformable continuum model that accounts for the physical properties of the membrane and the helix insertion to predict curvature sensing behavior and is in good agreement with existing experimental data. Specifically, we show that the insertion can be modeled as a local change to the membrane’s spontaneous curvature,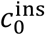. Using physically reasonable ranges of the membrane bending modulus к, and a 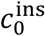 of ∼0.2-0.3 nm^-1^, this approach provides excellent agreement with the energetics extracted from experiment. For small vesicles with high curvature, the insertion lowers the membrane energy by relieving strain on a membrane that is far from its preferred curvature of zero. For larger vesicles with low curvature, however, the insertion has the inverse effect, de-stabilizing the membrane by introducing more strain. The membrane energy cannot be directly predicted analytically, due to shape changes from surface relaxation around the anisotropic insertion. We formulate here an empirical expression that captures numerically calculated membrane energies as a function of both basic membrane properties (bending modulus к and radius *R*) as well as stresses applied by the inserted helix (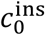 and area *A*_ins_). We show that the shape relaxation energy has a similar magnitude to the insertion energy, with a strong nonlinear dependence on 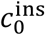. We therefore predict how these physical parameters will alter the energetics of helix binding to curved vesicles, which is an essential step in understanding their localization dynamics during membrane remodeling processes.

## I. Introduction

The recruitment of cytosolic proteins to membranes is an essential step in a variety of membrane remodeling processes, including clathrin-mediated endocytosis [1] and cell division [2, 3]. Proteins that participate in membrane remodeling contain membrane binding domains that use positively charged interfaces to specifically target negatively charged lipids such as PI(4,5)P_2_ on cell membranes [4, 5]. In addition to this electrostatic interaction, these proteins exploit additional mechanisms, including helix insertion, scaffolding, crowding, and entropy gain by disordered proteins [6-9]. These mechanisms cause proteins to bind more strongly to more highly curved membranes, driving them to both sense and induce membrane curvature [10]. Amphipathic *α*-helices are common protein domains that are frequently found in peripheral membrane proteins like septins, epsins, endophilins, and amphiphysins. Playing key remodeling roles in cell division and endocytosis, these proteins insert themselves into a single leaflet of a membrane, where they can sense curvature independently of any additional curvature sensing mechanisms [8]. The stronger binding of helix-containing domains to membranes of high curvature can thus control their localization dynamics, helping to regulate subsequent steps in assembly and remodeling. Understanding how the strength of membrane binding depends on helix insertion and membrane properties is thus an essential component of predicting the spatial control of protein localization and corresponding remodeling dynamics.

Several lines of experimental evidence support curvature sensing by amphipathic helices and its direct coupling to membrane deformations and membrane energy changes. Tethered vesicle assays visualize increased binding to highly curved membranes for domains with *α*-helix [8, 11]. Without the *α*-helix present, these same domains are insensitive to membrane curvature, demonstrating the significance of the helix in curvature sensing. The insertion or the *α*-helix changes the membrane local curvature and induces curvature generation [12-14]. Equilibrium observations of proteins bound to small unilamellar vesicles (SUVs) can be converted into dissociation constants by determining the relative partitioning of proteins to smaller vesicles (Fig 1a). These dissociation constants can then be used to extract changes of energy upon binding (Fig 1b). This data provides a basis for our modeling comparison, where we can then address how the key parameters of curvature, bending modulus, tension, insertion size, and spontaneous curvature control membrane energies following helix insertion.

**Figure 1.**
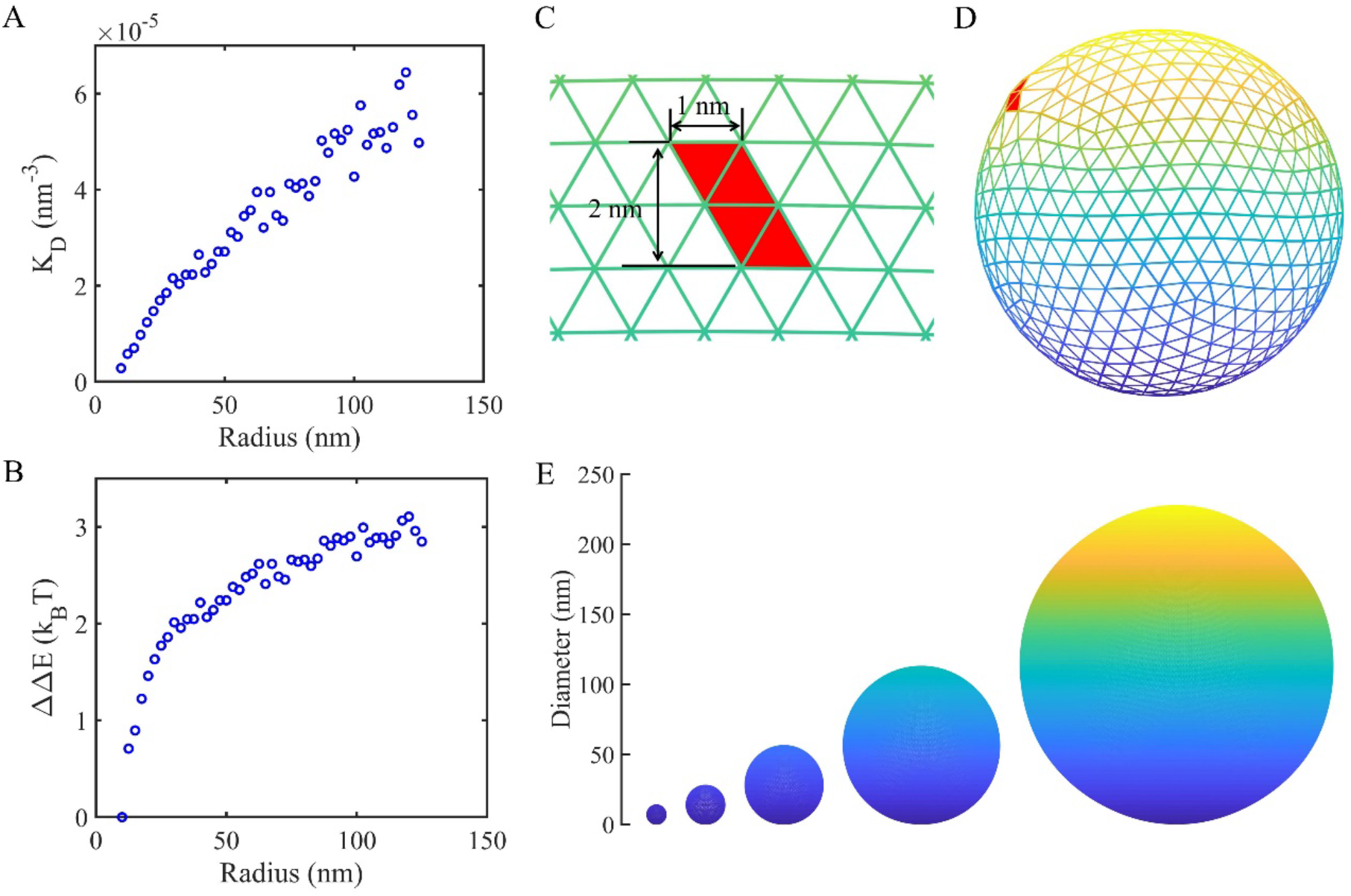
Experimental curvature sensing by ENTH domains with amphipathic helices and the corresponding model design to quantify these results. (a) The dissociation constant of the amphipathic helix-containing ENTH domain is stronger with more highly curved (smaller) vesicles. The solution concentration of the ENTH domain is 150 nM and the PI(4,5)P_2_ density on the vesicle is 0.125 nm^-2^. (b) From the K_D_, we can measure the difference in binding energy with vesicle size, using the smallest vesicle as the zero point. The positive energy changes reflect weaker binding. (c) The helix insertion is modeled as occupying four adjacent triangles on the vesicle surface (R=7nm), which are assigned a nonzero spontaneous curvature, 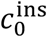, while the rest of the surface has a spontaneous curvature of zero. (d) The insertion modifies the membrane energy, with equilibrated structures deforming to produce local bulges that increase with larger 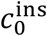. Here 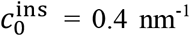 and R = 7 nm. (e) Five vesicles of different sizes in our simulations. The vesicle radii are 7 nm, 14 nm, 28 nm, 56 nm and 112 nm from left to right.

Modeling and simulation using other approaches have illuminated several ways in which helix insertion alters membrane structure and stress. Molecular dynamics simulations have measured depth and orientation of helices in bilayers, deformations of the surface and lipid structure [15, 16], and corresponding changes to membrane stress around the insertion [17]. However, these simulations are limited to relatively small (nanometer scale) and typically flat membranes. Curved all-atom membranes have shown that the underlying deformation matches molecular factors, like protein shape [18], but measurements of the energy changes arising from helices inserted into membranes of varying curvature have remained intractable using molecular dynamics. In contrast, models that use elasticity theory have quantified how stress profiles and energetics [19] in initially curved membranes will respond to helix insertion differentially, depending on how the curvature was generated and the depth of the insertion [20]. However, the modeled membrane patch varies with only two variables (thickness in one dimension, arc length in the other dimension), whereas our modeled vesicles exist in 3D space. These elasticity calculations thus only capture variation along one axes of principal curvature on a surface, assuming translational symmetry along the other. They cannot directly model spherical vesicles, where a highly localized and anisotropic insertion will impact curvature along both principal axes.

Deformable continuum membrane surfaces are an attractive model for studying membrane mechanics because they can adopt diverse geometries in 3D space, and their shape and energy will relax in response to perturbations. The calculations are relatively efficient and capture how the material bending modulus of the membrane, the membrane tension, and osmotic pressure will impact energetics and membrane shape [21]. Local perturbations driven by proteins adsorbed to the surface can be modeled via changes to the membrane’s spontaneous curvature *c*_0_, with values that can vary from 0 to ∼1 nm^-1^ [22]. The spontaneous curvature can vary spatially across the membrane surface, driving changes in membrane shape and tension [23], thus providing an effective material parameter that captures changes to membrane stress on the outer vs inner bilayer leaflets [22]. Continuum models can be coupled to models with attached and diffusing proteins [24-26] to capture interactions that drive membrane-mediated collective behavior [27], or response of cell-shape to flow[28]. Thus, these models provide a flexible platform for integrating mechanical responses with environmental changes or biochemical interactions. We apply this detailed curvature model here to protein curvature sensing, showing that changing the local spontaneous curvature is an effective parameter for capturing a helix insertion.

In this paper, we first describe the model design, and the approach used to compare model results with *in vitro* experiments measuring surface coverage on SUVs of varying curvature. We then show how the spontaneous curvature of the insertion, 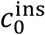, and the bending modulus к have dominant effects on the membrane energy changes following insertion. Both the size and spread of the insertion further modulate the magnitude of the energy changes. In contrast, constraints on the area and volume that would arise due to area compressibility and osmotic pressure have minimal impact on controlling energetic responses to insertion. With these results we are able to define parameter regimes that produce excellent agreement with the experimental observations of curvature sensing. We validate that the results are robust to our numerical methods, including mesh size, integration scheme, and optimization protocol. Finally, through our numerical results we derive an empirical formula that predicts how membrane energies will change following insertion as a function of variations in 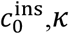, radius and insertion area. This expression can thus be used to estimate curvature sensing by amphipathic helices without additional computational measurements.

## II. MODEL DESIGN

### Continuum membrane model

The membrane is modeled using a continuum thin-film surface captured via a triangular mesh using the subdivision limit surface method [21] (Fig 1). The energy of the membrane is due to a bending energy (via the Helfrich Hamiltonian [29]) and constraints on the volume and area, defined as:

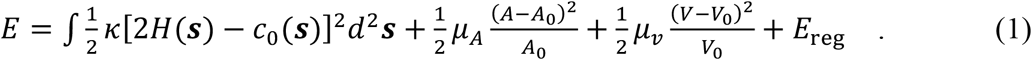

The first bending energy term integrates over all positions on the surface ***s***, and varies with mean curvature, *H(****s***), and the spontaneous curvature of the membrane, *c*_0_(***s***), where *κ* is the constant bending modulus. *H(****s***) is the mean of the calculated curvature along the two principal axes at point ***s***; for a sphere of radius *R*, it is a constant 1/*R* at all points. *H(****s***) thus changes when the membrane deforms. The next two terms capture the area and volume constraints, with respective coefficients *μ*_A_ and *μ*_v_. *A* is the membrane area, *A*_0_ is the target area of the membrane, *V* is the vesicle volume, and *V*_0_ is the target volume of the vesicle. The regularization energy, *E*_reg_, is added to eliminate the in-plane shearing deformations of the triangular mesh as the structure is optimized. This technical rather than physical constraint (due to the numerical mesh) should go to zero in equilibrated structures, and in Methods we describe specific forms we tested to minimize its contribution to the total energy.

Although the insertion is localized to a few mesh points, it drives the local region of the surface to bulge and deform to minimize the energy after the insertion, and thus requires integration over the 3D surface to evaluate energy changes (Methods). To study the curvature sensing effect, we ran simulations on vesicles with five different radii, as follows: R = 7 nm, 14 nm, 28 nm, 56 nm, 112 nm respectively, as shown in Fig 1e. The curvature is defined as the inverse of the vesicle radius, so the initial vesicles have curvature in a range of 0.009∼0.14 nm^-1^, which covers a range of relevant membrane curvatures in biological systems.

### Modeling helix insertion

The spontaneous curvature of bilayers is dependent on the lipid composition, and for the vesicles studied here, with symmetric leaflets, one can assume the spontaneous curvature is zero, meaning the membrane prefers to be everywhere flat. The insertion of the oblong α-helix into a leaflet of the bilayer will induce conformational changes of nearby lipids. We therefore model the effect of the insertion as a local change to the spontaneous curvature of the membrane [22]. The area of this local change is chosen to mimic the size of the α-helix domain, which occupies about 2 nm^2^, i.e. 1 nm in width and 2 nm in length, on the membrane [6, 30]. We thus select 4 triangles of the mesh to assign a nonzero value of c_0_, which we will refer to as 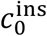 (Fig 1c). In the Results we verify that the conclusions are unchanged with either higher mesh resolution, or with a more diffuse spreading of the local change in 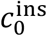. The rest of the membrane surface retains a spontaneous curvature of zero. Optimized structures thus can form a bulge around the insertion (Fig 1d). In this paper, we focus on the impact of one ENTH binding to the membrane, effectively assuming each binding event is local and independent of one another.

### Comparison to experimental observables

The experimentally measured coverage of ENTH proteins per vesicle of radius *R* is reported in Ref. [11], along with the experimental methods. We compute the corresponding *K*_D_ via *K*_0_ = [*P*]^eq^*ρ*_*l*_^eq^/*ρ*_*el*_^eq^, where [*P*]^eq^ is the concentration of ENTH in solution, *ρ*_2_^eq^ is the free lipid density on the surface, and *ρ*_*el*_ ^eq^ is the density of membrane-bound ENTH proteins. We exploit that both the proteins and the lipid sites are in great excess to the number of bound complexes, such that [*P*]^eq^ = [*P*]_tot_ (150nM) and 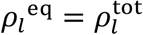. The lipid binding-site density is calculated from the 7.5% mol fraction of lipids that are PI(4,5)P_2_, as 0.125 copies/nm^2^ on all vesicles. The bound protein densities vary from 0.0038 to 0.0002 copies/nm^2^. This *K*_D_ is thus dependent on radius (Fig 1a), and the energetics of binding of one ENTH to the membrane surface can be extracted via the well-known relation

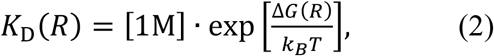

where ΔG is the free energy difference between an unbound ENTH and the membrane surface, relative to the bound state, and [1M] is the standard state free energy. For binding of ENTH domains with the helix removed, the *K*_0_ becomes effectively independent of radius (Fig S1).

Now we assume that this binding free energy can be broken into two contributions: a chemical potential which is independent of membrane curvature, due to electrostatic interactions between protein and lipids (−*μ*), and a mechanical energy due to helix insertion into the bilayer (ΔE). Without the helix, the ENTH domain still binds the membrane, but the binding is insensitive to curvature (Fig S1), hence a curvature independent term. Thus Δ*G*(*R*) = −*μ* + Δ*E*(*R*) and inserting this into Eq 2, we can write

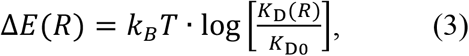

where *K*_D0_ is independent of the membrane shape. To compare with simulations, we measure how these mechanical energies change relative to that of the smallest vesicle (Δ*E*_ref_), defining:

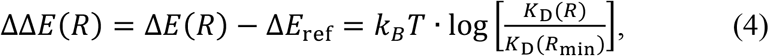

where *R*_min_ = 10 nm (Fig. 1b). When the proteins and lipid binding sites are in excess of the membrane bound proteins (true here), this expression is independent of [*P*]_tot_ and 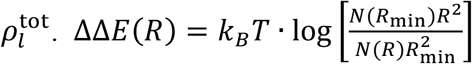, where *N* is the copies of membrane bound proteins per vesicle of size *R*.

## III. RESULTS

### IIIA Spontaneous curvature of the insertion drives opposite changes in energy in small vs large vesicles

The impact of our helix insertion on the membrane energy is controlled by its size (2nm^2^) and by its spontaneous curvature, 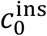. For 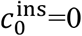, the insertion would not change the membrane energy at all for any vesicle size, which we measure via

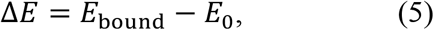

where *E*_bound_is the membrane energy with one helix bound and *E*_0_ = 8*π*к [29] is the energy of the spherical vesicle with no insertions. This unperturbed energy *E*_0_ results from the cost of bending the membrane into a sphere, when it prefers a flat curvature (*c*_0_ across the unperturbed surface is zero), and is independent of vesicle radius.

As we increase 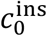 of the insertion from 0.01 nm^-1^ to 0.3 nm^-1^, we find that for smaller vesicles, the insertion lowers the cost of bending the membrane, producing ΔE < 0 (Fig 2a). For these highly curved vesicles, the insertion thus improves the stability of the bound system. Conversely, for larger vesicles, we see the opposite effect. Once R>28nm, insertions with increasing 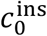 cause an increase in the cost of bending the membrane (ΔE > 0), thus de-stabilizing these flatter surfaces.

**Figure 2.**
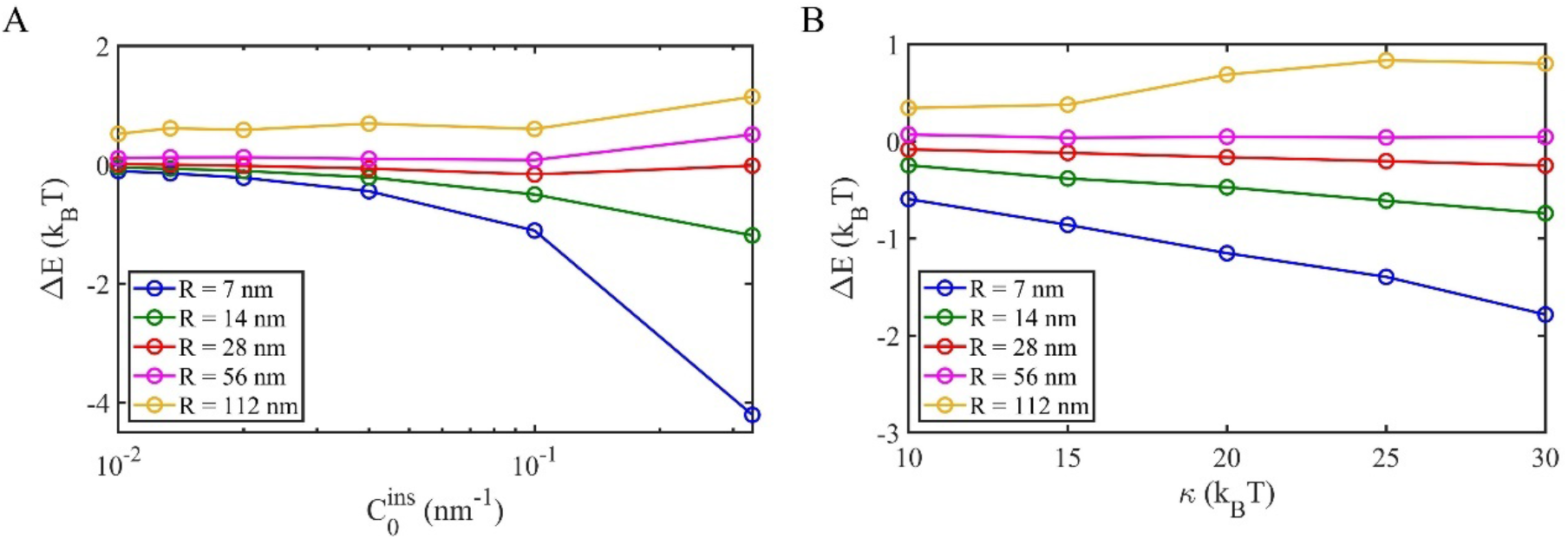
The addition of insertions to spherical vesicles can stabilize or de-stabilize the bending energy dependent on vesicle size and bending modulus. (a) As 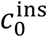 increases, the insertion has a larger impact on the membrane energy, stabilizing small vesicles and de-stabilizing large ones. *κ* =20 k_B_T. (b) As the bending modulus *κ* increases, the insertion is again more stabilizing for small vesicles, but de-stabilizing for large vesicles. 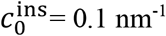.

These inverse costs and benefits of adding the insertion result from the initial strain that the membranes are under by being forced into curved (instead of flat) enclosed vesicles. For the highly curved small vesicles, the local mean curvature *H(****s***) is high and introducing an insertion that prefers higher curvature relieves strain in the membrane, even as it drives local shape changes. With larger vesicles, the local mean curvature *H(****s***) decreases, and the creation of local shape changes around the insertion eventually costs energy. The transition occurs at ∼R=28nm, where the corresponding membrane curvature is about 0.036 nm^-1^. The transition between the positive and negative energy responses cannot be predicted from the vesicle size and insertion spontaneous curvature alone, as it depends on the relaxation of the surface around the insertion, as described further in Section IIIG.

### IIIB. Stiffer membranes also show opposite response to insertion in small vs large vesicles

The response of the membrane energy to changes in bending modulus κ is also coupled with the vesicle size. For small curved vesicles, as we stiffen the membrane against bending (larger κ) the insertion produces a greater benefit in stabilizing the membrane (Fig 2b), retaining a ΔE < 0 and 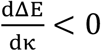. This result is again in agreement with the insertion relieving the strain in the membrane, which increases in the unperturbed vesicle with larger κ. For intermediate vesicles, the energy change becomes less sensitive to changes in κ, until we reach R=112nm. Now we see the opposite trend, where the increasing stiffness causes a larger cost to adding the insertion, producing 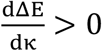. The sign of ΔE does not change with κ, only the magnitude, as we quantify in Section IIIG. However, we note that in Fig 2b, the spontaneous curvature of the insertion zone is fixed at 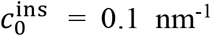 for each data point, which means we effectively assume that the insertion (e.g. the insertion depth) is not influenced by the stiffness of the membrane. In reality, the insertion parameters of a specific *α*-helix type could be coupled to the membrane stiffness. So, given a specific helical structure, the response of the membrane energy to varying bending modulus *κ* could reflect simultaneous changes in both *κ* and 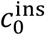. A relationship between *κ* and 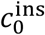 would depend on molecular properties of the protein-lipid interactions and is beyond the scope of this paper.

### IIIC. Area and volume changes have minimal impact on curvature sensing

The membrane energy is dependent on changes in volume and surface area due to insertion (Eq 1), although we find they make negligible contributions relative to the bending energy. The volume constraint reflects an influence of osmotic pressure, where water may pass in and out of the vesicle to change its volume. The coefficient *μ*_*v*_ controls the penalty to changes in volume, and we find that over a broad range of values, it has minimal impact on the membrane energy after insertion (Fig S2). Similarly, the expansion or compression of the membrane area upon insertion is controlled by the coefficient *μ*_A_, where experiments estimate this membrane elastic modulus in the range of 230 – 260 pN/nm [31]. Here again, over a broad range of values of *μ*_A_, we see minimal changes to the membrane energy, indicating that global area changes upon insertion are not significant contributors to curvature sensing (Fig S2). In all of our simulations (unless otherwise noted), we thus use fixed values of *μ*_v_ = 83.4 pN/nm^2^ and *μ*_*A*_ = 250 pN/nm.

### IIID. Helix insertion energies are sensitive to local curvature, not membrane surface area

Our results above show that the effect of the insertion on membrane energy is sensitive to the curvature of the membrane surface, as it changes with vesicle size. For a helix insertion with a fixed value of 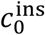, we can thus see that as the vesicle gets smaller and more curved, the insertion drives more stable energies 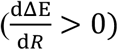, with a steeper benefit occurring for larger values of 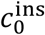 (Fig 3b). Because both the curvature and the membrane surface area are changing with vesicle size, we further tested whether this curvature sensing would be retained on a fixed surface area. We thus generated a single enclosed red-blood-cell-shaped membrane (by decreasing the vesicle volume [32]) that exhibits variations of curvature across its surface. Four different points with different curvatures on the surface were selected as the insertion zone (Fig 3c). Our simulation results recapitulate the same curvature sensing phenomena, where binding to the most highly curved region produces the largest benefit in membrane energy changes, and binding to regions of negative curvature produces a cost in membrane energy (Fig 3d). The curvature sensing ability is robustly driven by the local curvature and resulting deformation around the insertion, and not just the relative size of the perturbation to the total surface area.

**Figure 3.**
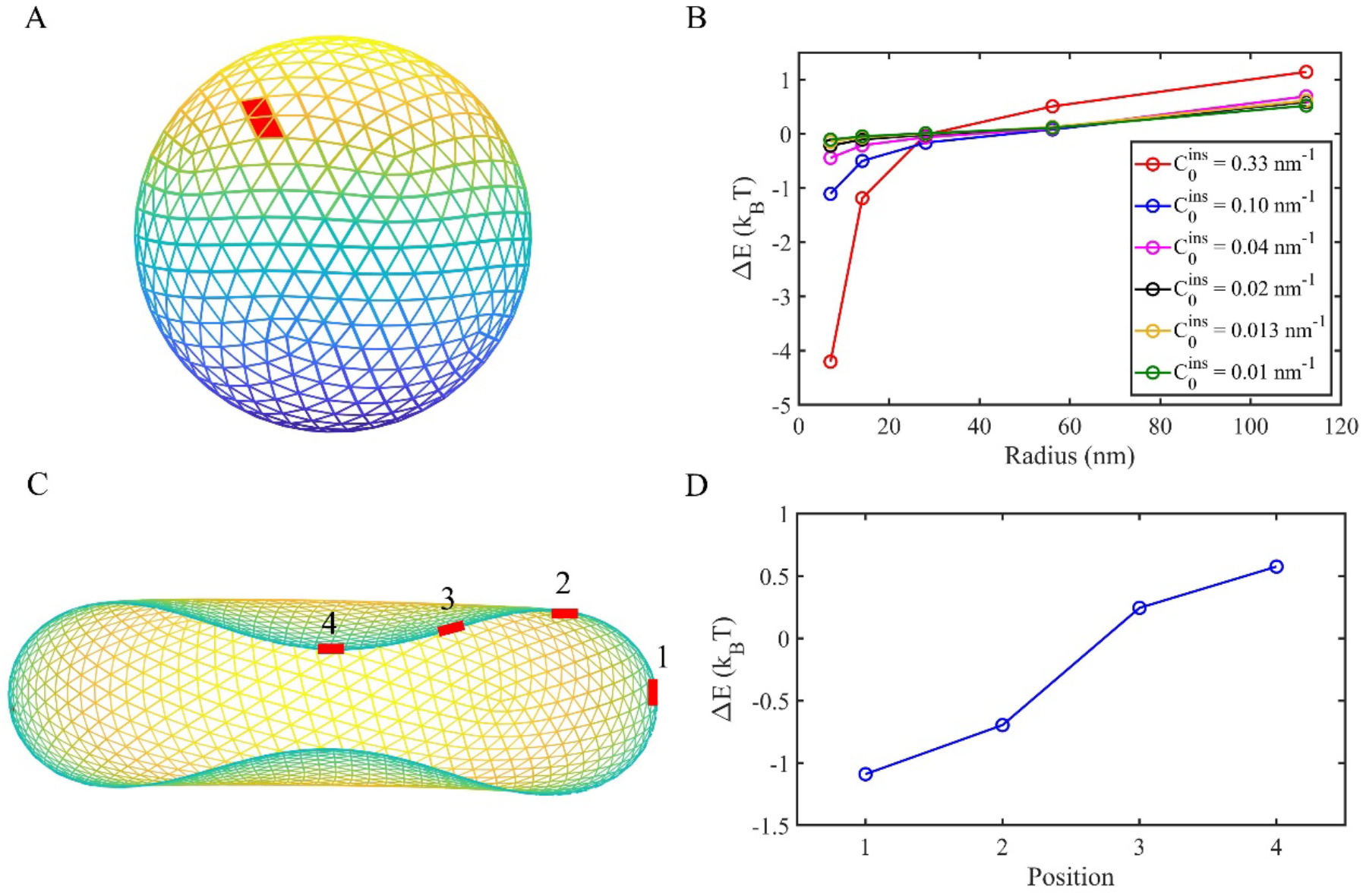
Helix insertions will drive stronger binding to more highly curved membranes. (a-b) Curvature sensing on five spherical vesicles of different sizes. (a) Spherical vesicle R = 7 nm and the insertion position (red color). (b) Insertions increase the stability of the membrane energy (ΔE < 0) more robustly with smaller radius and correspondingly higher curvature (1/R). The magnitude of this effect depends on the spontaneous curvature of the insertion 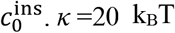. (c-d) Curvature sensing on one ‘vesicle’ with heterogeneous curvatures. (c) Side view of oblate structure. To get this asymmetric oblate structure, we started with the spherical vesicle R = 14 nm and set the target volume V_0_= 0.65×4πR^3^/3, target area S_0_ = 4πR^2^. A much stronger area and volume constraints were added with μ_v_ = 8.34×10^4^ pN/nm^2^ and μ_A_ = 2.50×10^4^ pN/nm. κ = 20 k_B_T and 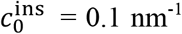. The membrane curvature at the positions labeled 1,2,3,4 is 0.14, 0.08, 0.06, −0.06 nm^-1^ respectively. (d) The membrane energy change ΔE is more stabilizing with higher curvature.

### IIIE. The model recapitulates in vitro measurements of curvature sensing

To most directly compare the simulation results to the experimental results, we measure the energy change upon insertion relative to the value in the smallest vesicle (Δ*E*_ref_ (R=7nm)), so following Eq. 4,

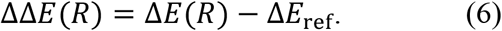

We see excellent agreement between the shape of the energetic changes between both our numerical results and the experiment (Fig 4), where in fact more than one set of 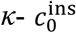 values agree quantitatively with the experiment. Hence, within physically reasonable values of the membrane bending moduli, a softer membrane reproduces the data with a larger spontaneous curvature 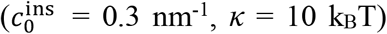 or a stiffer membrane matches with a weaker spontaneous curvature 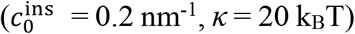. We can see then that if the spontaneous curvature is either too large 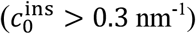 or too small 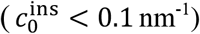, then the numerical results cannot reproduce observed bending energy changes for reasonable value of *κ*.

**Figure 4.**
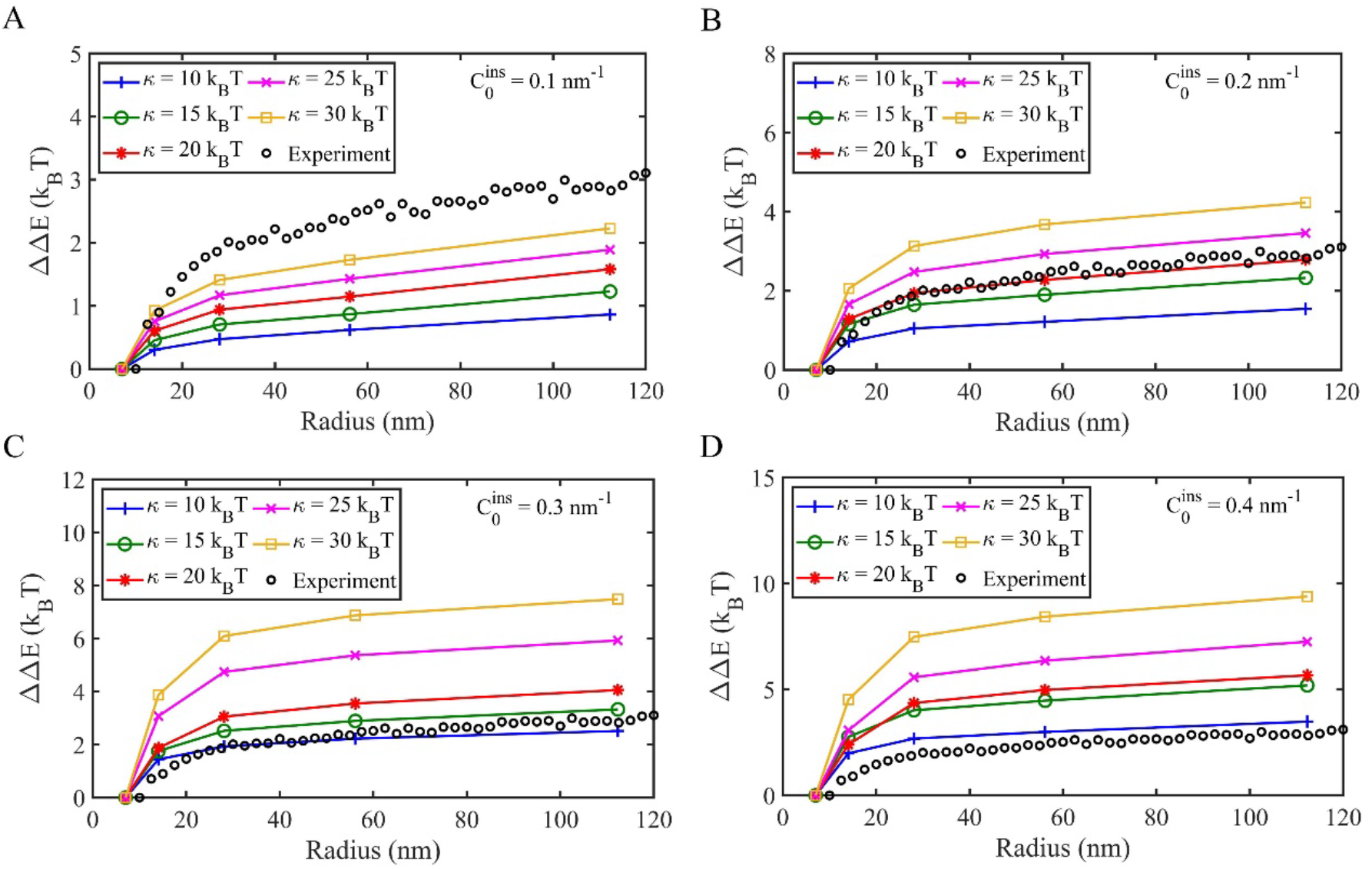
Numerical simulations can reproduce the *in vitro* curvature sensing results for realistic bending moduli and spontaneous curvature values. In all panels, the experimental data is shown in black open circles. a) For a lower spontaneous curvature 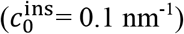, the curvature sensing is too weak to reproduce the experiment. b-c) For values of 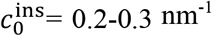, we see excellent agreement with the experimental energetics, where the agreement is dependent on pairing of 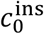 and *κ*. d) With large spontaneous curvature 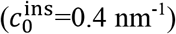, the energetic changes are much larger than are observed in experiment.

### IIIF. Curvature sensing is retained when the insertion area is spread out

In the above models, our insertion locally modified the spontaneous curvature, and immediately dropped off in adjacent surface elements. To test the effect of having a longer distance impact on neighboring membrane, due to a stressed distribution of lipids around the insertion [17], we expanded the region of non-zero spontaneous curvature around the insertion (Fig 5a). The value of 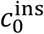 is thus largest at the center of the insertion, and decays with distance away, which we model using a Gaussian function as:

**Figure 5.**
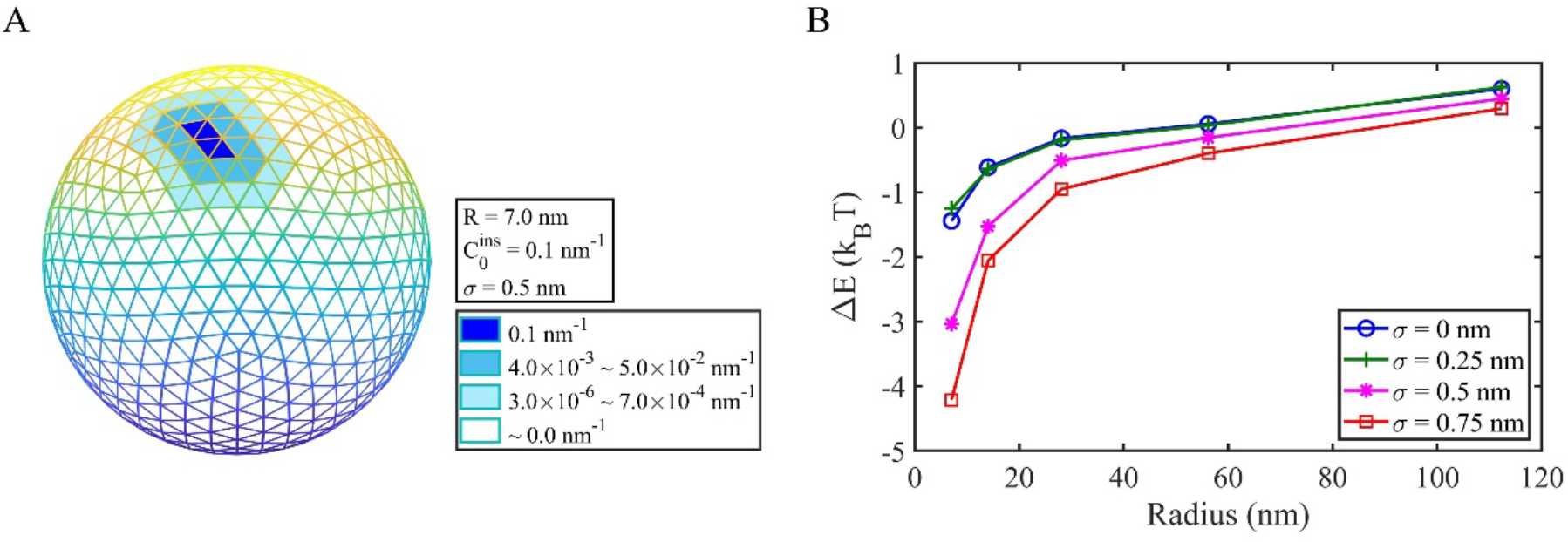
Expanding the insertion area amplifies the changes in membrane energy. (a) The insertion can be modeled as impacting the spontaneous curvature of neighboring membrane, in a distance-dependent way. (b) As the spread of the insertion region is increased (increased *σ*), we see larger responses in the membrane energy change. 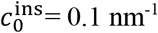 and *κ* =20 k_B_T.

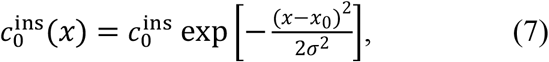

where *x* is the distance to the insertion zone center *x*_0_, and σ is the width of the spread. The induced stress or strain is limited to ∼1 nm scale around the insertion [17], so we choose 2σ ≤1.5 nm. We find that the curvature sensing effect is robustly retained, with a larger spreading of the insertion producing a larger change in the membrane energy (Fig 5b).

### IIIG Predictive model for membrane energy changes following insertion captures effect of membrane shape changes

To combine all of our numerical results into a simpler mathematical framework, we derived a phenomenological expression to predict how the membrane energy changes would vary with 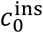, *κ*, vesicle radius R, and insertion area *A*_ins_. This expression clarifies how the observed energy change results not only from the local change in spontaneous curvature at the insertion (which can be calculated analytically), but the membrane shape changes that occur following membrane relaxation around the insertion.

We write the total observed membrane energy change as

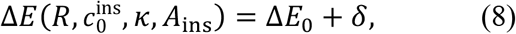

where Δ*E*_0_ is the change in bending energy due solely to changes in spontaneous curvature at the insertion. Therefore, *δ* captures energy changes due to relaxation of the membrane shape to relieve induced strain in the surface. To calculate Δ*E*_0_ we can use Eq. 1, where we have the membrane bending energy of the area *A*_ins_ on the unperturbed spherical vesicle is 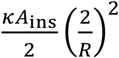, with the spontaneous curvature of the membrane being 0. Once the amphipathic helix inserts into this area *A*_ins_, the bending energy on this area is 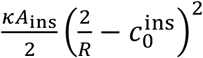 with 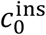 being the effective spontaneous curvature on this binding area *A*_ins_. We thus have:

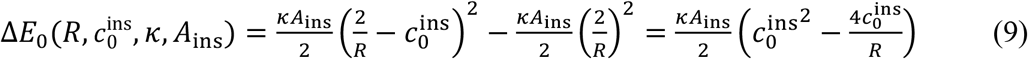

as the membrane energy change from one insertion before the relaxation of the stress or strain (Fig 6a).

**Figure 6.**
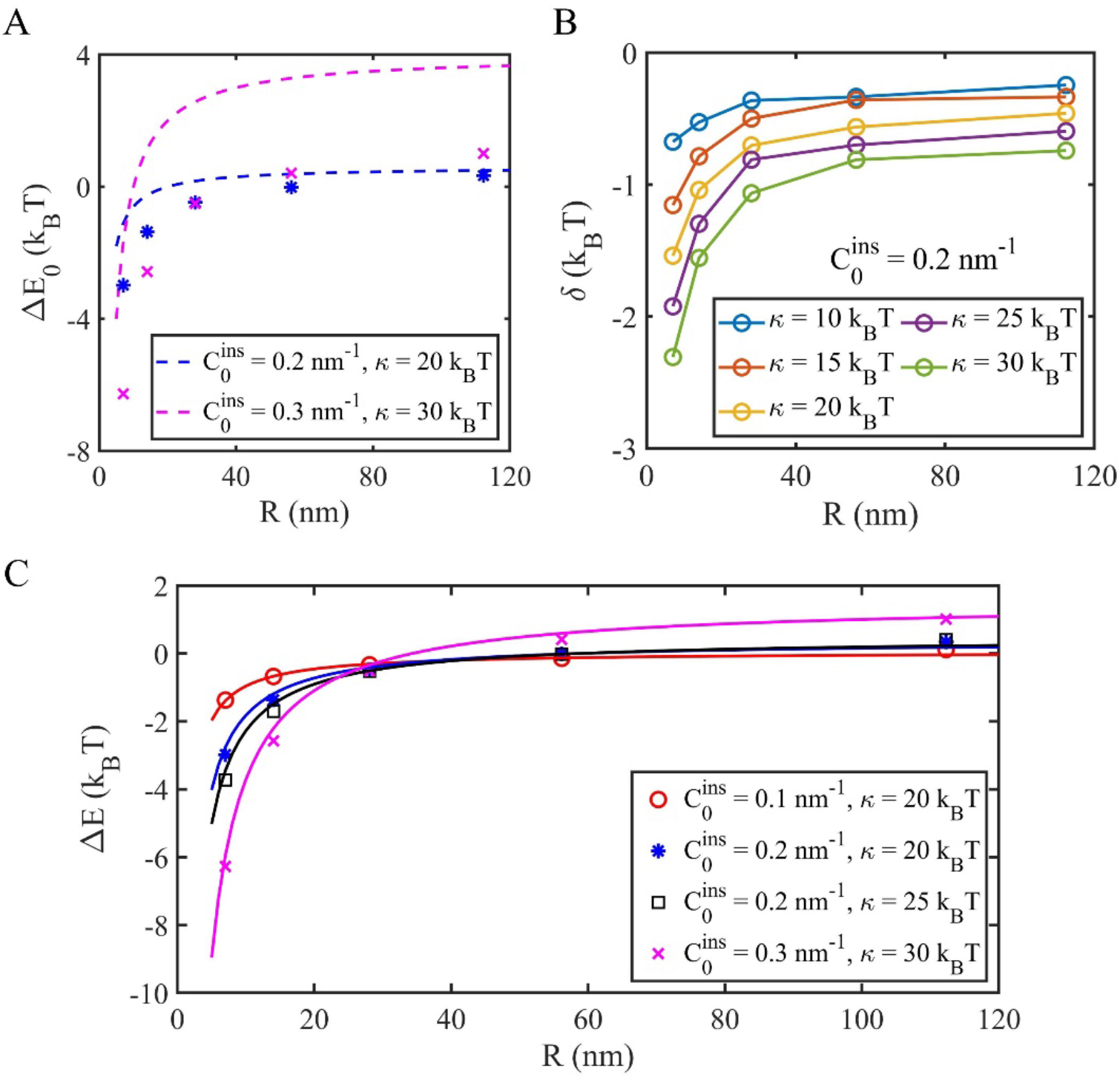
Empirical formula predicts dependence of membrane energy change on physical parameters due to the insertion and subsequent membrane shape changes. A) The energy change Δ*E*_0_ due only to insertion with nonzero spontaneous curvature (before shape relaxation) significantly overestimates the energies observed at equilibrium. Dashed lines are analytical expression of Eq (9), and data is the total energy change drawn the same color and symbols as in part (C). B) Energy change *δ* due to membrane shape relaxation, calculated from the numerical simulation energies and Eq (8). C) The total energy change can be fitted well by Eq (11).

We use our numerical results to derive an expression for *δ*. Our results show δ is always a negative value (Fig 6b), which means the relaxation process always causes the membrane energy to decrease, whereas Δ*E*_0_ is positive when 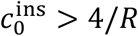. From our analysis, we find that δ varies with all four parameters, 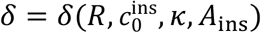. The simulation results show δ ∝ *κ* and that δ is a linear function of *R*^−1^ (Fig S3), which is the same dependence that Δ*E*_0_ has (Eq 9). For the insertion parameters, however, we find that *δ* has distinct scaling with 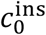 and *A*_ins_. Specifically, we find that 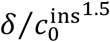 is independent of 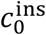, and *δ*/*A*_ins_^1.25^ is independent of *A*_ins_ (Fig S3). By fitting our numerical data, we thus recover a final practical expression of:

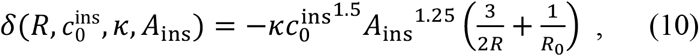

where R_0_ = 10 nm is a fit parameter necessary to capture the apparent plateauing of *δ* at negative values when R→∞.

By combining Eqs (8-10), we have the analytical expression of the membrane energy change due to one helix insertion as:

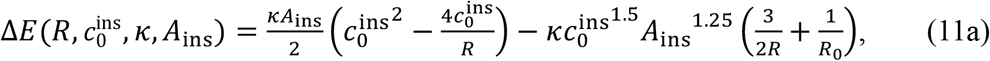

or

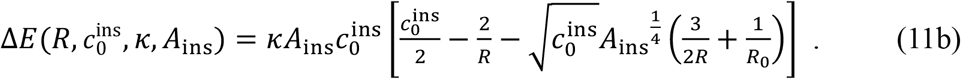

Eq (11) provides excellent agreement with the numerical data as shown in Fig 6c, recovering the proper limits that as the size or spontaneous curvature of the insertion goes to zero, there is no change in the membrane energy, as expected. This model further predicts when the helix insertion will cause stabilization (Δ*E* <0) or destabilization (Δ*E* > 0) to the membrane energy, dependent on 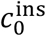, *R*, and now also *A*_ins_. The sign is thus independent of к, as seen in Fig 2b. This expression shows that the membrane energy changes for vesicles is most sensitive to changes in 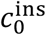, which is coupled to the vesicle radius most strongly via the membrane shape changes, as seen in the last term of Eq. 11a 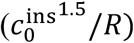. The trend is similar for the insertion size, where it couples more strongly to the vesicle radius in the membrane shape changes (*A*_ins_^1.25^/*R*). The magnitude of the energy change following relaxation is comparable to the energy change due to the insertion (Fig. S3e), meaning that the contribution of the membrane shape changes in response to helix insertion cannot be ignored when quantifying the strength of helix localization to membranes.

## IV. Discussion

Curvature sensing by amphipathic helices emerges from their localized disruption of, primarily, the leaflet of the bilayer where they embed. The energy change that results from this localized perturbation is up to a few k_B_T, based on experimental measurements. We show here that curvature sensing by amphipathic helices and the corresponding energy changes can be accurately captured by deformable continuum membrane models, despite lacking an explicit double leaflet structure. Instead, the spontaneous curvature of an insertion area, which is a material property reflective of stresses induced on only one leaflet, can effectively couple inserted helices to the membrane bending energy. Our numerical results predict stronger binding of helices to membranes of higher curvature. The form of the curvature-dependent binding energy is excellent quantitative agreement to experiments. Furthermore, using literature standard values for к (15-20 k_B_T)[33], and predicted values for 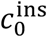 (0.2-0.5 nm^-1^)[34], the experimental observations are directly within the range of reasonable parameters. The energy change that accompanies helix insertion is due to the bending energy, and we decompose this energy change into two parts: the cost of the helix insertion (change in 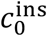) and the energy of the shape change following insertion. Both components make comparable contributions to the overall change in bending energy. We develop an empirical formula that can then predict these energy changes, which quite accurately captures dependence on the bending modulus, sphere radius, helix insertion size, and insertion spontaneous curvature. We can therefore predict when helix insertion acts to relieve stress in the membrane (highly curved vesicles) or introduce new strain (low curved vesicles). We verify that the observed energy changes are due to sensing of the local curvature around the insertion, as the result is retained in non-spherical surfaces of constant surface area, and as the helix insertion is spread.

We assume the binding of each amphipathic helix is independent of each other. This applies for the low concentrations used in the experiments here, where the density of proteins on the membrane surface never surpassed 0.0038/nm^2^ (< 5% surface coverage-Fig S4) and no clustering of proteins was observed [11]. However, at higher densities, the local shape changes could alter the binding energetics of subsequent proteins, leading to mechanically induced feedback. Mechanical feedback can alter rates of binding to membranes[35]. The spatial distribution or interactions between proteins on the membrane can also vary due to localized changes to bending energy and membrane shape[36]. At coverage above 20%, additional curvature induction mechanisms such as crowding [6], would enhance shape changes beyond helix insertion alone. The modeling approach used here is capable of quantifying even small changes in energy that could emerge due to cooperative effects. In future work we will address how feedback and cooperativity can drive enhanced or depressed recruitment to surfaces of varying curvature.

A limitation of the thin-film surface model is that it does not explicitly capture the thickness of the bilayer. The model thus cannot quantify how the stress profile in the membrane varies[37] from the embedded leaflet, where the helix causes stretching, relative to the presumably more compressed opposite leaflet. Where finer-grained detail is required, applications of material-elastic theory [37] to membrane patches thus have an advantage in this regard. Our approach here benefits from previous material-elastic studies that have predicted ranges of 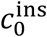 from helix shape and mechanical strain[34, 38]. Furthermore, membrane thickness has been shown to be an important variable for interactions between fully transmembrane proteins that span both leaflets [36, 39]. Studying the role of membrane thickness in continuum models can be achieved through coupling of two layers together, and hybrid methods that combine continuum membranes with atomistic proteins capture realistic deformations around transmembrane proteins [40, 41]. Here we found that capturing explicit membrane thickness was not necessary, as the spontaneous curvature accurately quantified helix-induced membrane strain.

Overall, the modeling approach used here offers an accurate, experimentally verified platform to study membrane shape changes and bending energies arising from adsorbed proteins. To quantify curvature sensing by amphipathic helices, we found it to be efficient across multiple changes to material properties of the membrane and the insertion, even over ∼k_B_T or smaller energy changes. The direct comparison between quantitative experiments and modeling provides a mechanism to determine coarse material parameters of proteins, where here we found that inserted helices have a spontaneous curvature of 0.2-0.3nm^-1^. Modeling membranes at the mesoscale has proved critical for studying key steps in processes from clathrin-mediated endocytosis [42], to fluctuations in red-blood cell membranes[43], where molecular approaches are simply intractable. Coupling deformable membranes to dynamics using reaction-diffusion methods [44] offers a mechanism to capture the added complexity arising from mechanical feedback or reversible protein-protein interactions, with broad applications in membrane remodeling in the cell [45]. Our code is therefore provided open-source under a Gnu Public Licence (GPL) at github.com/mjohn218/NERDSS/continuum_membrane.

## V. Methods

### VA Set-up of vesicles

An enclosed spherical triangular mesh is set up by the Loop’s subdivision scheme at the radius of interest [46]. The limited surface area is calculated as the vesicle area S, and the volume enclosed by the vesicle area is calculated as the vesicle volume V.

### VB Energy minimization

The equilibrium state of the vesicle is produced by minimizing the total energy using nonlinear conjugate gradient methods (NCG) [47]. The force on each vertex is expressed as the derivative of the total energy to the vertex position

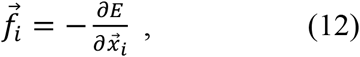

where 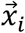 and 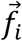 are the position and nodal force on vertex *i* respectively, and the detailed expression can be found in [21, 32]. The total energy includes terms due to the regularization and area constraint on the insertion described below. As criteria for stopping the minimization (finding the optimum), we use a mean nodal force is smaller than 10^−2^ pN and that the energy curve slope is (*E*_*i*+500_ − *E*_*i*_)/500 < 10^−3^ (E_*i*_ is the total energy in simulation step *i*), as shown in Fig 7.

**Figure 7.**
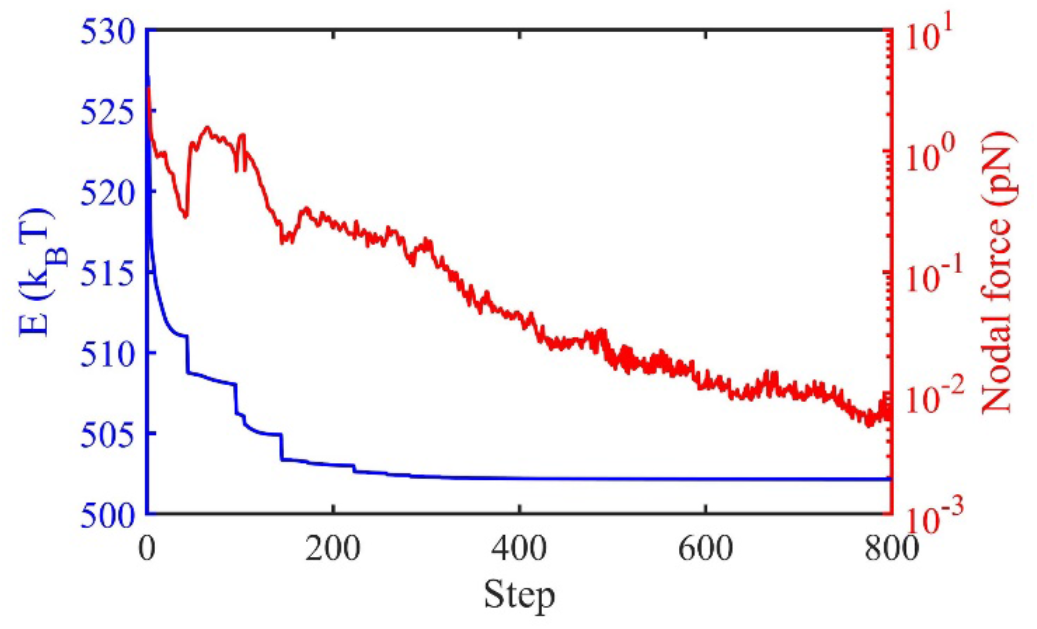
Energy minimization of the continuum membrane model following insertion. Simulation results from a vesicle with R = 14 nm and one insertion on the surface, 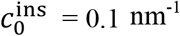, *κ* =20 k_B_T.

### VC Insertion Area constraint

We constrain the area of the insertion zone, as the nonzero 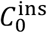 makes the triangular mesh nonuniform around the insertion, and we do not want the mesh deformation to change the area of the insertion (which is typically fixed at 2 nm^2^). Therefore, we tried two methods which produce very similar results, and neither of which measurably impacts the total energy of the system, which is dominated by the bending energy (Fig S5). Using an edge length energy we have:

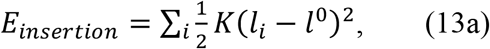

where *l*_*i*_ is the edge length of the insertion zone, 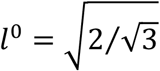 nm is the targeted length for the insertion zone, and *K* is the spring coefficient. The sum of Eq (13a) covers all the edges of the insertion zone. Alternatively, the insertion area can be constrained via a local area constraint:

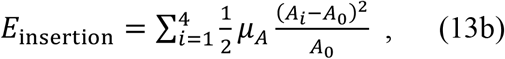

where *A*_*i*_ is the area of one triangle in the insertion zone, A_0_ = 0.5 nm^2^ is the target area of the insertion zone triangle (the total area of the insertion zone is 2 nm^2^, and the insertion zone has four triangles so each triangle should have area 0.5 nm^2^), and *μ*_*A*_ is the membrane area elasticity modulus as in Eq (1). To use Eq (13b) for the insertion area constraint, we need to separate out the insertion area from the global area constraint in Eq (1).

### VD Evaluation of surface integrals

The numerical solution of the integral over the surface in this theoretical model (Eq 1) is calculated by second order Gauss-quadrature. We validate that the second order of Gauss-quadrature is sufficient to produce converged energy estimates, and the higher-order and more expensive quadrature schemes are not necessary (Fig S6). We also validate that the fineness of the triangular mesh doesn’t influence the energy calculation (Fig S7), verifying that the resolution used here is sufficient to accurately measure energy changes following insertion.

### VE Regularization Energy

To eliminate the in-plane shearing deformations of the triangular mesh, we add a regularization energy [32]. The regularization energy has two forms depending on whether the triangular element is too biased from the equilateral shape. The function to describe the shape of triangular element *i* is defined by

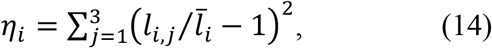

where *l*_*i,j*_ is the edge length and 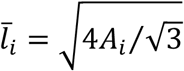 is the target edge length with *A*_*i*_ being the triangular element area. A large *η*_*i*_ means the triangle is more deformed, and here in our simulations we use *η*_0_ = 0.2 as the criteria determining whether the triangle shape is too deformed. If *η*_*i*_ > *η*_0_, the regularization energy for this triangular element *i* is

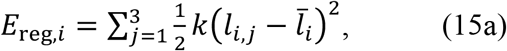

where *k* is the coefficient of this spring-type energy. If *η*_*i*_ ≤ *η*_0_, the regularization energy for this triangular element *i* is

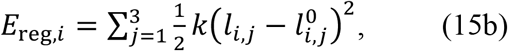

where 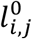 is the edge length we choose to use which is called the reference structure. Then the total regularization energy is the sum of all the *N* triangular elements of the mesh

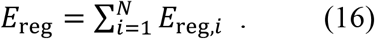

The regularization energy controls sizes of mesh elements which improves numerical integration over the surface, and thus is a technical constraint on the numerical method rather than physical constraint on the membrane energy, so it should converge to 0 when the system reaches the equilibrium state. The reference structure (value of 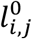) needs to be updated as the simulation evolves, and we update it when the energy optimization slows. This update method ensures that Eq (16) will converge to 0 and that the regularization works effectively on remeshing the triangular mesh [32]. Note that Eq (15) is not a continuous function, so it may cause a problem to find an efficient step size during the NCG energy minimization, but practically this problem can be solved by restarting the simulation or by shutting down Eq (15a) for several simulation steps.

## Acknowledgements

MEJ gratefully acknowledge funding from an NIH MIRA R35GM133644. We acknowledge supercomputing resources provided by ARCH at Johns Hopkins University, and the NSF-MRI funded rockfish cluster. We thank Dr Alexander Sodt and Prof Brian Camley for helpful discussions.

