## Supplemental Information for "A continuum membrane model predicts curvature sensing by helix insertion"

**Supporting Figures:**

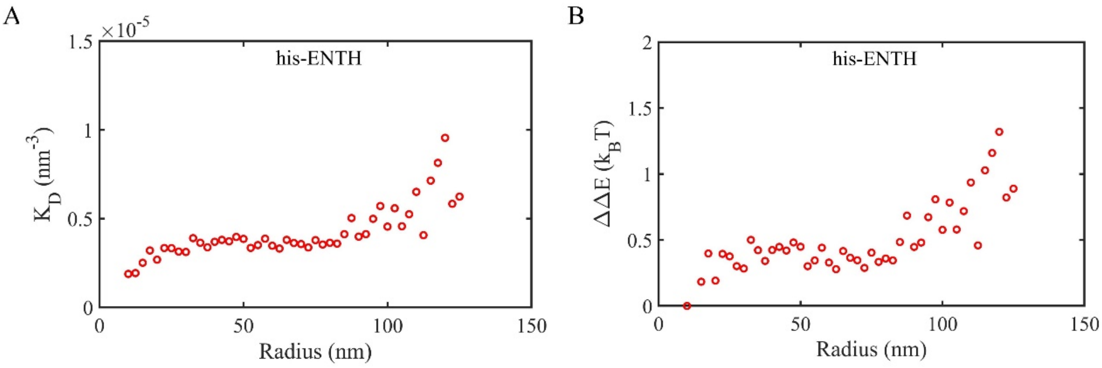


**Figure S1. Binding of ENTH without its amphipathic helix present is insensitive to curvature.** (a) Experimental surface coverage measurements of the hexa-histidine tag ENTH (his-ENTH) that lacks the amphipathic helices are converted to dissociation constants as described in the main text. (b) The corresponding membrane energy change is nearly flat, due to the curvature insensitivity across all vesicle sizes. The solution concentration of the his-ENTH experiment is 25 nM.

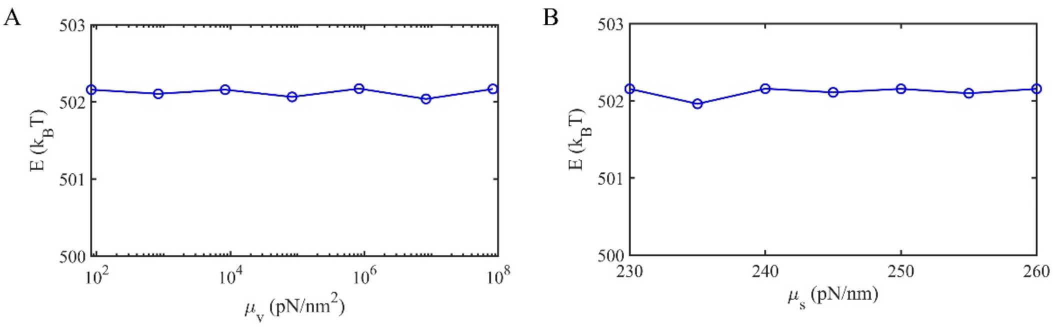


**Figure S2. Influence of the osmotic pressure and the membrane area elasticity on membrane energy changes.** (a) Changes to the volume constraint coefficient, $\mu_v$, reflecting changes in osmotic pressure, do not cause discernable changes in the membrane energy following insertion. $\mu_A$ = 250 pN/nm. (b) Changes to the area elasticity $\mu_A$ similarly do not impact the membrane energy change following insertion. $\mu_v$ = 83.4 pN/nm$^2$. The size of the vesicle is R = 14 nm, the insertion size is 2nm$^2$ (Fig 1), with $c_0^{\text{ins}}$ =0.1 nm$^{-1}$.

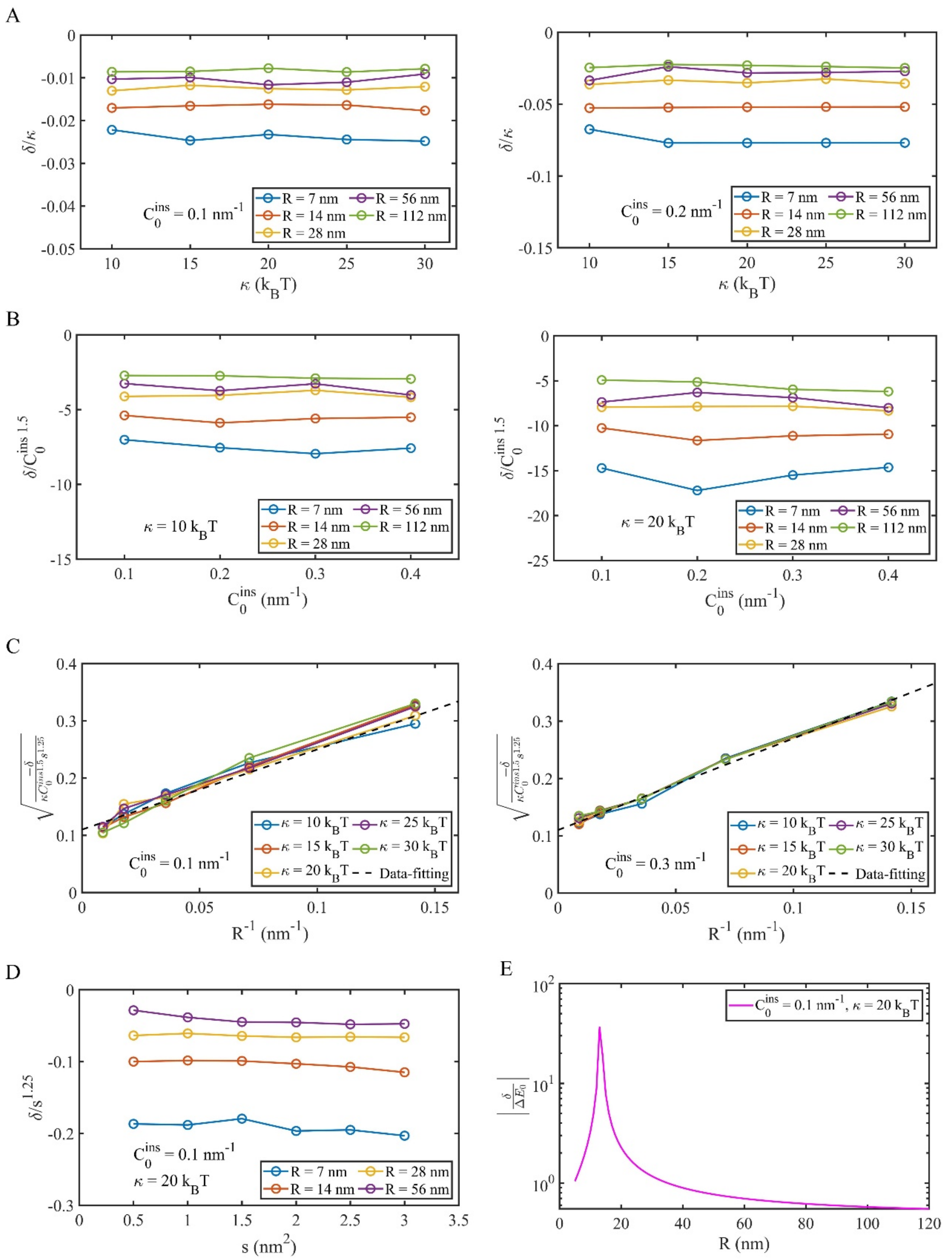


**Figure S3. Dependence of the membrane shape change energy, δ, with variations to membrane and insertion parameters.** (a) δ ∝ $\kappa$. (b) δ ∝ $\left(c_0^{\text{ins}}\right)^{1.5}$. (c) δ ∝ $R^{-1}$. (d) δ ∝ $A_{\text{ins}}{}^{1.25}$. (e) The magnitude of δ is comparable to the energy change due to the insertion, making a larger contribution for small-intermediate vesicles of a curvature similar to the size of $c_0^{\text{ins}}$.

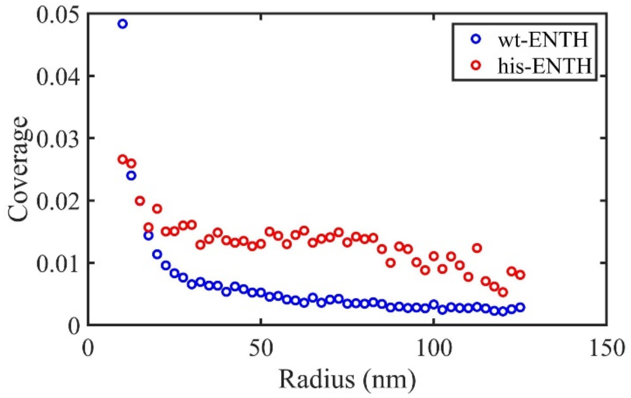


**Figure S4. Experimentally measured surface coverage of ENTH on vesicles.** The coverage is defined as $N\pi r^2/(4\pi R^2)$, where N is the copy number of ENTH bound on the vesicle, $r$ = 2 nm is the ENTH size and R is the vesicle radius.

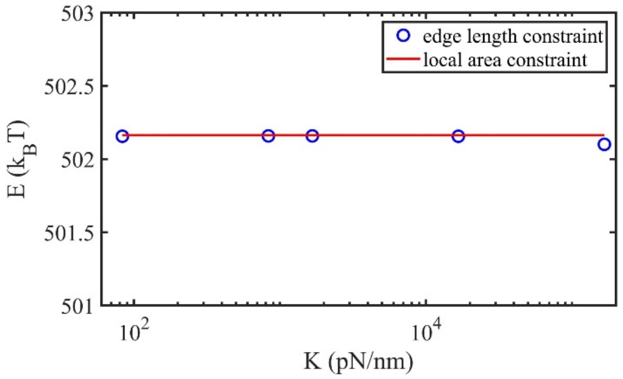


**Figure S5. Membrane energy following insertion is not sensitive to constraint choice on insertion region** For the edge length constraint method, we tried five different values of K. For the local area constraint method, we used $\mu_A$ = 250 pN/nm. Simulations were carried out with one insertion with $c_0^{\text{ins}}$ = 0.1 nm$^{-1}$, $\kappa$ = 20 $k_B$T, $\mu_v$ = 83.4 pN/nm$^2$.

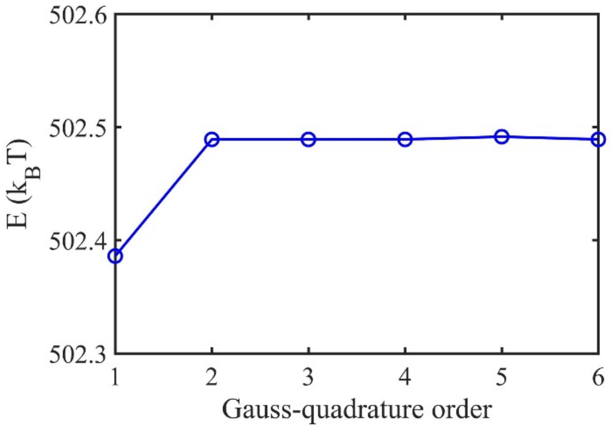


**Figure S6. Second-order Gauss-quadrature is sufficient to accurately integrate**

**bending energies.** The simulation is carried out on the vesicle of R=28 nm with $c_0^{\mathrm{ins}}$= 0.1 nm$^{-1}$, $\kappa$ = 20 $k_BT$, $\mu_v$ = 83.4 pN/nm$^2$ and $\mu_A$ = 250 pN/nm.

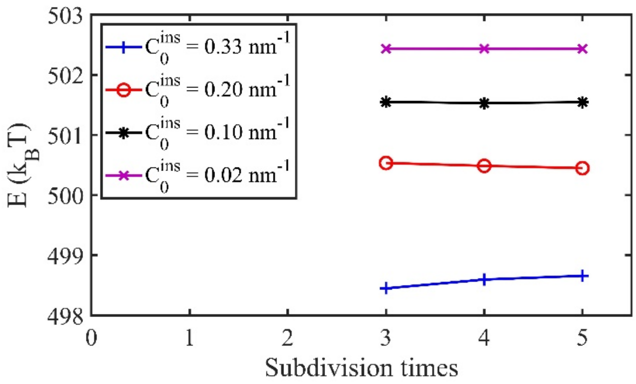


**Figure S7. Energy calculations following insertion are not sensitive to the fineness of the triangular mesh beyond 3 subdivisions.** The triangular mesh is generated by means of Loop's subdivision scheme. The more subdivision times give the finer mesh. Simulations here were carried out with the vesicle R = 7 nm and one insertion bound.
